# Differential expression of a disease-associated *MRE11* variant reveals distinct phenotypic outcomes

**DOI:** 10.1101/2025.07.15.664809

**Authors:** McKenna B. DeFoer, Ahmed M. Mostafa, Andrea J. Hartlerode, Steven K. Orban, Keegan McDonough, Sophie Quirk, Brianna K.L. Ferguson, David O. Ferguson, JoAnn M. Sekiguchi

## Abstract

The MRE11 DNA nuclease plays central roles in the repair of DNA double-strand breaks (DSBs) as a core component of the heterotrimeric MRE11/RAD50/NBS1 (MRN) complex. MRN localizes to chromosomal DSBs and recruits and activates the apical DSB repair protein kinase, ATM, which phosphorylates downstream substrates to elicit cellular DNA damage responses. Pathogenic variants in *MRE11* cause the genome instability disorder ataxia-telangiectasia-like disorder (ATLD). The first ATLD patient allele identified, *ATLD1*, is a nonsense mutation that deletes 76 amino acids from the MRE11 C-terminus and results in markedly reduced levels of MRE11-ATLD1 and the entire MRN complex. This region of the C-terminus has been demonstrated to function in DNA binding, mediate functional protein interactions, and undergo post-translational modifications that regulate MRE11 nucleolytic activities. We previously demonstrated that transgenic mice expressing low wildtype MRN exhibit severe phenotypes, including small body size, anemia, and cellular DNA DSB repair defects. Thus, it is currently unknown whether reduced MRE11-ATLD1 and MRN levels, loss of the C-terminus, or both cause disease-associated phenotypes. In this study, we generated transgenic mouse models that express near endogenous or significantly reduced levels of MRE11-ATLD1 to determine the *in vivo* importance of the MRE11 C-terminus. We observe that reduced MRE11-ATLD1 expression leads to anemia, bone marrow failure, extramedullary hematopoiesis, and impaired lymphocyte development, similar to mice expressing low wildtype MRE11. In contrast, higher expression of MRE11-ATLD1 results in a subset of moderate phenotypes, indicating that loss of C-terminus has limited impact on MRN functions *in vivo*. These findings have implications for clinical predictions of ATLD patients harboring pathogenic *MRE11* variants that impair MRE11 function and/or impact MRN protein levels.

## Introduction

Pathogenic germline variants in DNA damage response and repair genes underlie genome instability disorders characterized by a wide spectrum of disease sequelae, including bone marrow failure, immunodeficiency, developmental defects, neurodegeneration, and cancer predisposition. The constellation and severity of phenotypes observed in individual patients are highly variable and influenced by both the mutated gene and specific pathogenic variant(s). Ataxia-telangiectasia-like disorder (ATLD, MIM 604391) is a genome instability syndrome caused by biallelic germline variants in the *MRE11* gene (1). *MRE11* encodes a DNA repair nuclease that is a core component of the MRE11/RAD50/NBS1 (MRN) complex, which plays critical roles in sensing and repairing DNA double-strand breaks (DSBs). The MRN complex binds DSB ends and recruits the ATM protein kinase to sites of damage where it is activated.

Activated ATM phosphorylates a multitude of protein substrates to initiate the DNA damage response, eliciting cell cycle arrest, DNA repair, apoptosis, or senescence (2). MRN also stabilizes DNA ends in close proximity and facilitates long-distance DNA tethering through the long coiled-coil domains of RAD50 (3–5). Furthermore, MRE11 possesses both exo- and endonuclease activities that process DSB ends to enable repair by homologous recombination (HR), nonhomologous end-joining (NHEJ), or microhomology mediated end-joining (MMEJ) (6–10). Thus, MRN plays critical roles in both initiating the DNA damage response and directly repairing broken DNA ends.

ATLD patients typically present with progressive cerebellar ataxia and oculomotor apraxia, similar to the related disorder ataxia-telangiectasia (AT, MIM 208900) caused by pathogenic variants in the <u>A-T</u> <u>m</u>utated (*ATM*) gene. However, some ATLD patients also exhibit cognitive deficiencies, microcephaly, developmental defects, myoclonic ataxia, and cancer predisposition, with considerable variation in severity and age of onset (11–15). Disease-associated *MRE11* mutations span the entire gene and include missense, nonsense, and splice site variants that cause exon skipping (12). Many of these mutations result in markedly reduced MRE11 protein levels and consequently the entire MRN complex (1, 11, 16–18). This reduction confounds a clear understanding of whether ATLD phenotypes arise from specific defects in MRE11 function, reduced MRN levels, or both.

The first ATLD patient allele identified, *ATLD1*, is a nonsense mutation that truncates the C-terminal 76 amino acids from the 708aa human MRE11 protein (p.R633*, referred to as MRE11-ATLD1 herein) (1) (Fig. 1A) and significantly reduces MRN complex levels in both patient-derived cells and a gene-targeted mouse model (1, 19). It is notable that the C-terminal region deleted in ATLD1 patients has distinct functions. Previous studies demonstrated that the MRE11 C-terminus binds to the cyclin-dependent kinase, CDK2, and this interaction is necessary for efficient phosphorylation and activation of CtIP, a cell cycle regulated DSB repair factor (9). Phosphorylated CtIP interacts with MRN and activates MRE11 endo- and exonucleolytic activities that are required for DNA end resection (20–22). Structural and biochemical analyses have revealed that the C-terminal region interacts with intact double-stranded DNA via an ATP-independent mechanism distinct from its DNA end-binding activity (5). This interaction is hypothesized to facilitate genome scanning (5) and/or DSB-independent activities of MRN, such as maintaining the integrity of highly active transcription units (23). Additionally, several amino acids residing in the MRE11 C-terminus undergo DNA damage-induced post-translational modifications, including phosphorylation and lactylation, which were demonstrated to regulate MRE11 nucleolytic activities (24, 25). Thus, a growing body of evidence suggests the MRE11 C-terminus plays multiple, important roles in the DNA repair functions of MRN. However, its *in vivo* functions remain undefined.

**Figure 1.**
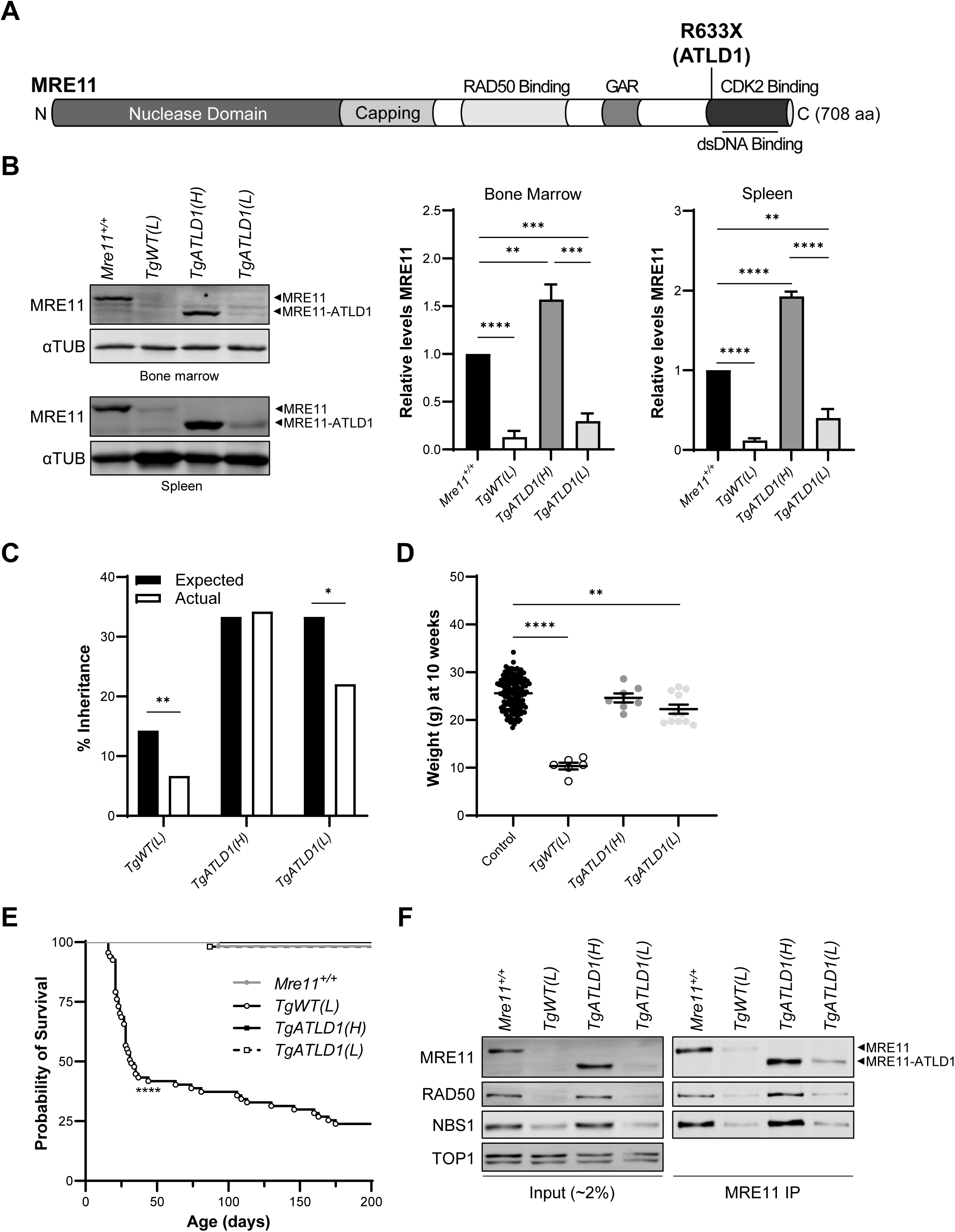
Low MRE11 and MRE11-ATLD1 levels cause sub-Mendelian birth rates and growth deficiency. A) Schematic of the domain structure of human MRE11. Location of the R633X, *MRE11-ATLD1* mutation, within the C-terminal region is indicated. B) Western blots and quantitation of the MRE11 levels in mouse organs from young adult mice (5-13 weeks). α-Tubulin is the loading control. Mean and SEM are shown relative to *Mre11^+/+^* (n ≥ 3 mice). C) Percentage inheritance from matings involving *Mre11^+/−^* x *Mre11^−/−^TgMre11* (or *Mre11^+/−^TgMre11* for *TgWT(L)* matings). *n* ≥ 120 offspring per cross. *Χ^2^≤0.05 and **Χ^2^≤0.01 indicate a significant difference between the actual and expected birth rates. No blinding, randomization, or power calculation for sample size was performed. D) Mouse weights from individual mice at ∼10 weeks of age. The control group contains all littermates that express *Mre11* from the endogenous locus (*Mre11^+/+^* or *Mre11^+/−^*) with or without a transgene. E) Kaplan-Meier survival analyses of transgenic mouse lines showing percent survival versus age (in days). Control (*n*=59), *TgATLD1(H)* (*n*=50), and *TgATLD1(L)* (*n*=53) survive longer than *TgWT(L)* mice (*n*=67). \*\*\*\**P*≤0.0001 by Log-rank test. F) MRN complex stability was determined via immunoprecipitation with an anti-MRE11 antibody and immunoblots were performed with antibodies to MRN components. For all panels, significance was determined via unpaired *t*-test (\*\**P*≤0.01, \*\*\**P*≤0.001, and \*\*\*\**P*≤0.0001).

To examine the *in vivo* importance of the MRE11 C-terminus, we have developed novel transgenic mouse models expressing MRE11-ATLD1 at near wildtype levels of endogenous MRE11 in an *Mre11^−/−^* background. To distinguish the impact of the *MRE11-ATLD1* allele versus reduced MRN levels, we generated transgenic mice expressing wildtype MRE11 (26) or MRE11-ATLD1 at significantly reduced levels. Unexpectedly, we found that mice expressing near endogenous levels of MRE11-ATLD1 do not exhibit severe organismal or cellular phenotypes. However, loss of the MRE11 C-terminus alone impairs B cell development at specific stages and results in elevated levels of chromosomal anomalies in unperturbed primary cells and irradiated murine embryonic fibroblasts. Thus, we demonstrate that the MRE11 C-terminus does have *in vivo* functions in facilitating efficient repair of chromosomal damage. In contrast, reduced expression of wildtype MRE11 or MRE11-ATLD1 results in marked and distinct developmental, hematopoietic, lymphopoietic, and cellular phenotypes. Overall, our study reveals that differential expression of MRE11-ATLD1 and wildtype MRE11 cause distinct phenotypic outcomes. These findings have important implications for predicting the clinical presentation and severity of phenotypes in ATLD patients.

## Results

### Generation of *Mre11-ATLD1* transgenic mice

To differentiate the impact of the *MRE11*-*ATLD1* disease allele and loss of the MRE11 C-terminus independent of reduced MRN complex levels, we generated a transgenic mouse model expressing the murine *Mre11-ATLD1* cDNA (nt1-1896 of the 2118nt full-length murine *Mre11*) from the well-characterized pCAGEN mammalian expression vector (Fig. 1A). We obtained 16 *Mre11-ATLD1* transgenic founders, which were crossed to *Mre11^+/−^* mice to subsequently generate lines expressing the transgene in an *Mre11^−/−^* background. Mice heterozygous for the *Mre11-ATLD1* transgene were used for all analyses. Importantly, mice expressing the transgene in an *Mre11^+/−^*background did not exhibit phenotypes in any analyses, thereby indicating the phenotypes observed in transgenic mice and cells were fully complemented by one wildtype *Mre11* allele.

Expression levels of the C-terminally truncated MRE11-ATLD1 protein were assessed in multiple tissues by western blotting. We identified transgenic lines that expressed MRE11-ATLD1 at levels similar to or higher than endogenous wildtype MRE11 (approx. 1.5-2-fold; Fig. 1B and data not shown), which we refer to as *TgATLD1(H)*. Concurrently, we obtained transgenic mice expressing MRE11-ATLD1 at significantly reduced levels (approx. 30-40% of endogenous MRE11), similar to ATLD1 patients and an *Mre11-ATLD1* knock-in mouse model (1, 19) (Fig. 1B). We refer to these mice as *TgATLD1(L)*. *TgATLD1(H)* mice are viable, born in Mendelian ratios, produce offspring, and are of similar size and weight as wildtype controls (Fig. 1C-E). In contrast, the *TgATLD1(L)* mice are born in sub-Mendelian ratios and are significantly smaller in size and weight; however, they do not exhibit shorter lifespans compared to controls (Fig. 1C-E). These phenotypes are less severe compared to transgenic mice expressing low levels of full-length *Mre11* (*TgWT(L)*), which exhibit sub-Mendelian live births, markedly lower body weights, and significantly reduced survival (Fig. 1C-E; (26)). We note that *TgWT(L)* mice express MRE11 and the MRN complex at approximately 10-15% of endogenous levels, which may result in more severe phenotypes compared to *TgATLD1(L)* mice that express relatively higher MRN levels (Fig. 1B, 1F). Nonetheless, these findings demonstrate that expression of MRE11-ATLD1 at near endogenous levels supports normal embryonic development, body weight, fertility, and lifespan.

Protein levels of the MRN subunits, RAD50 and NBS1, were assessed in *TgATLD1(H)* and *TgATLD1(L)* murine embryonic fibroblasts (MEFs) by western blotting (Fig. 1F). We detected reduced levels of the entire MRN complex in *TgATLD1(L)* cells, as previously observed in ATLD1 patient cells and gene-targeted mice (1, 19). However, RAD50 and NBS1 levels were unaffected in *TgATLD1(H)* cells. Furthermore, MRE11-ATLD1 expressed from the transgene appears to maintain interactions with RAD50 and NBS1 in MEFs, as determined by co-immunoprecipitation analyses (Fig. 1F).

### Reduced expression of MRE11 and MRE11-ATLD1 leads to defects in hematopoiesis

*TgATLD1(L)* mice are born with marked pallor (Fig. 2A), similar to newborn *TgWT(L)* mice that exhibit severe anemia (26). Complete blood counts (CBCs) in *TgATLD1(L)* mice revealed a significant decrease in red blood cells (RBCs), hemoglobin, and hematocrit with a concomitant increase in mean corpuscular volume (MCV), indicating severe macrocytic anemia as previously observed in *TgWT(L)* mice (Fig. 2B, C; (26)). In contrast, *TgATLD1(H)* pups are not pale, and CBC analyses revealed no significant differences in RBCs, hemoglobin, hematocrit, or MCV compared to wildtype controls (Fig. 2B, C). In addition to anemia, mice expressing reduced MRE11 or MRE11-ATLD1 exhibited reduced white blood cells, bone marrow cells, and thymocytes compared to wildtype controls (Fig. 2D; (26)). Interestingly, elevated platelet counts were also observed in *TgWT(L)* and *TgATLD1(L)* (Fig. 2C), a phenotype distinct from anemias arising in other genome instability syndromes such as Nijmegen breakage syndrome (caused by *NBS1* variants) and Fanconi anemia, which are characterized by abnormally low platelet counts (27–29).

**Figure 2.**
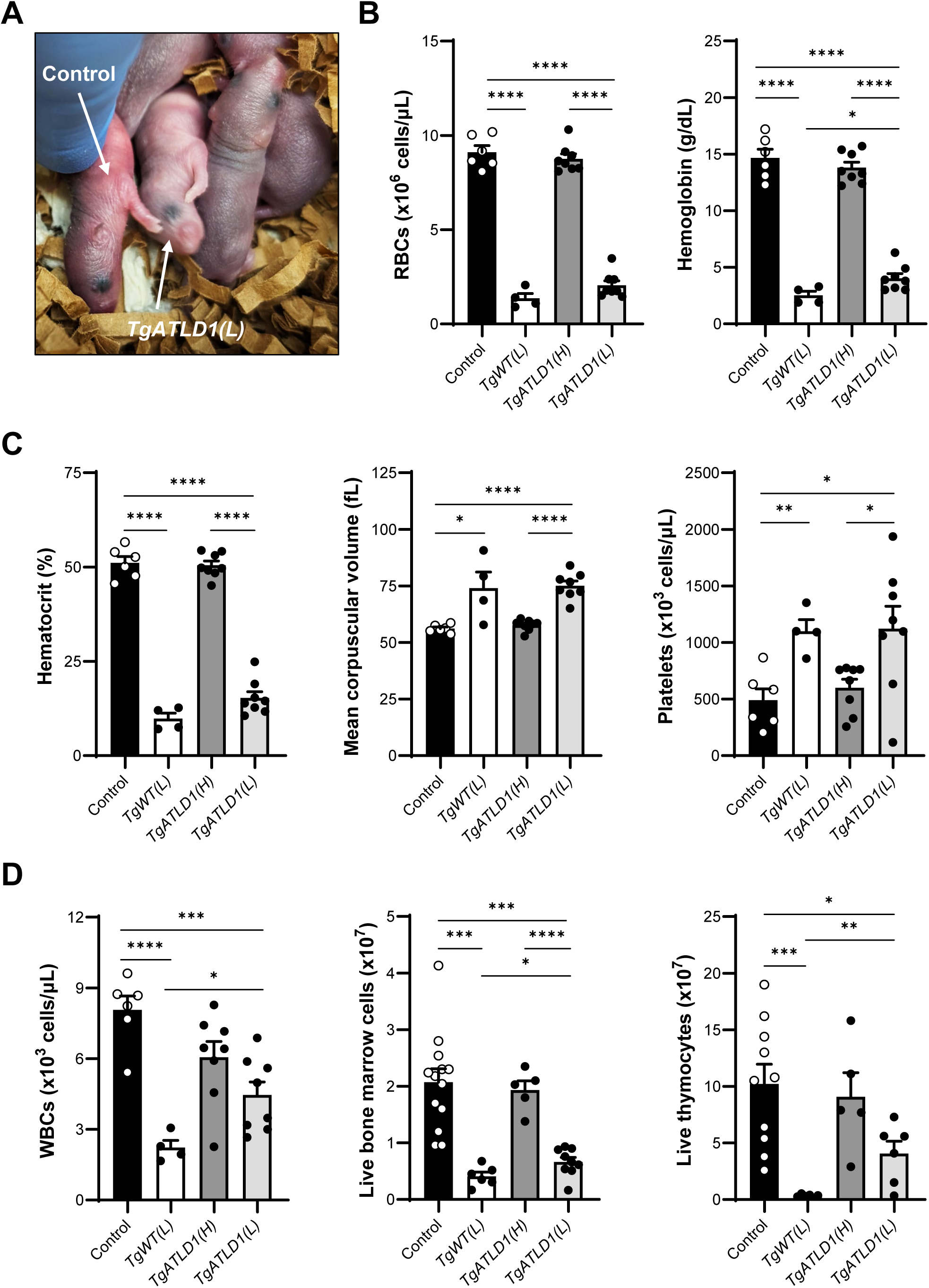
*TgATLD1(L)* mice exhibit severe anemia and signs of defective hematopoiesis. A) *TgATLD1(L)* pups are pale from birth compared to littermate controls that express normal levels of MRE11 (either endogenous *Mre11^+/+^* or *Mre11^+/−^*). Image is taken at postnatal day 2. B-D) Complete blood cell counts were performed on mice aged 28-100 days. (B) Red blood cell (RBC) and hemoglobin counts, (C) hematocrit, mean corpuscular volume, and platelet counts, (D) white blood cell (WBC) counts (left). D) Total number of live bone marrow cells (middle) and thymocytes (right) from *n* ≥ 5 mice per genotype. For all panels, each point represents an individual mouse, *n* ≥ 4 mice per genotype, mean ± SEM are plotted, and significance was determined via unpaired *t*-test (\**P*≤0.05, \*\**P*≤0.01, \*\*\**P*≤0.001, and \*\*\*\**P*≤0.0001).

To assess the underlying causes of the observed blood cell phenotypes, we examined hematopoiesis in the bone marrow by flow cytometry (Fig. 3A). *TgWT(L)* and *TgATLD1(L)* bone marrow exhibited a higher percentage of uncommitted, lineage negative (Lin^−^) hematopoietic cells and a concomitant decrease in mature effector cells (Fig. 3B). Further analyses of the hematopoietic stem and progenitor (HSPC) populations within the Lin^−^ populations revealed a significant increase in hematopoietic stem cells (CD150^+^CD48^−^c-Kit^+^Sca1^+^; HSCs) and common lymphoid progenitors (Lin^−^Sca-1^+^c-Kit^+^IL7R-α^+^; CLPs) in *TgWT(L)*, but not *TgATLD1(L)*, bone marrow (Fig. 3C). Populations of LS^−^K cells, common myeloid (CMP), granulocyte-monocyte (GMP), and megakaryocyte-erythroid progenitors (MEP) were more variable in *TgWT(L)* and *TgATLD1(L)* bone marrow, and most differences did not reach statistical significance compared to controls (Suppl. Fig. S1). In contrast, we observed no anemia or hematopoietic defects in *TgATLD1(H)* mice.

**Figure 3.**
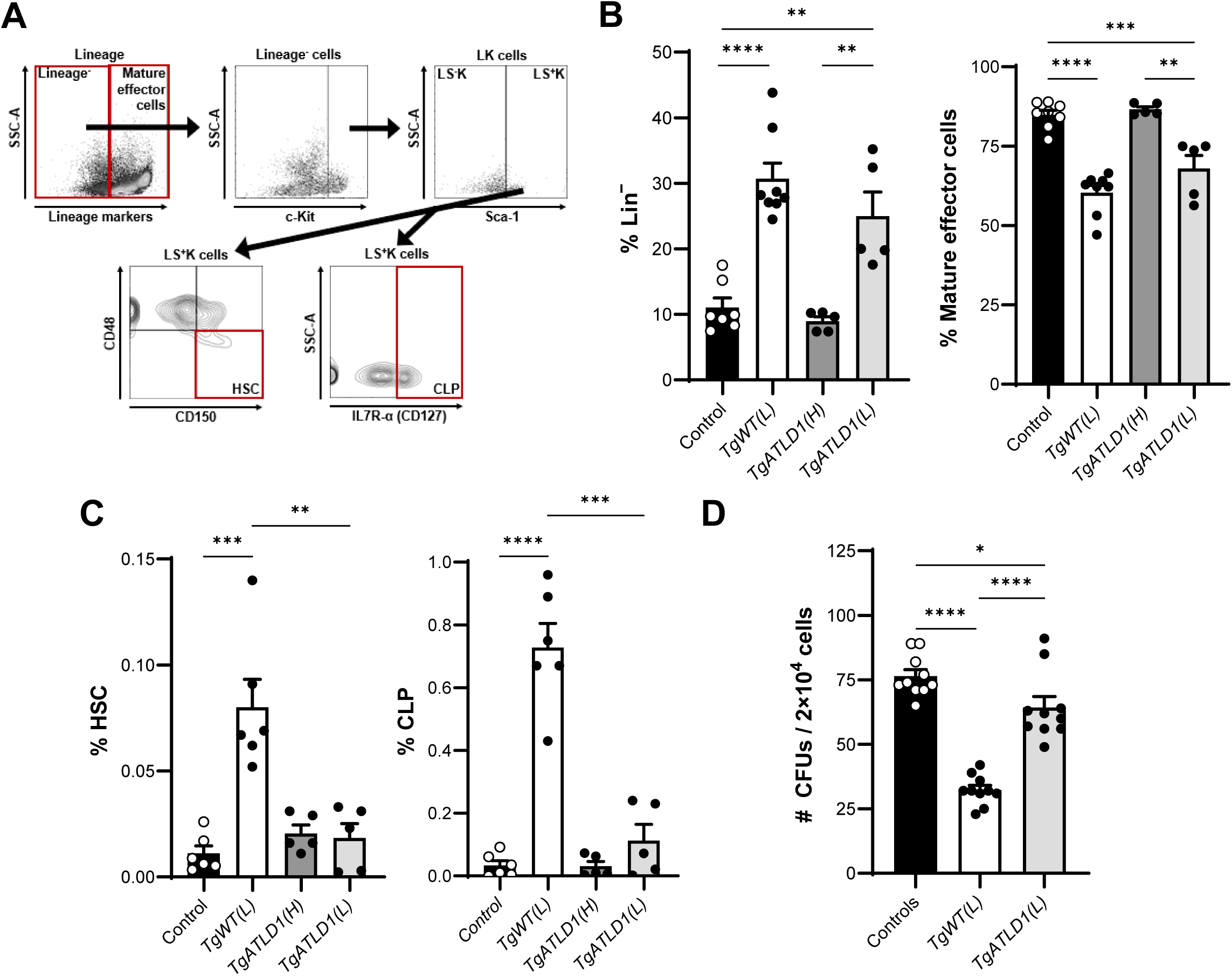
Reduced MRN levels impair differentiation and proliferation of hematopoietic progenitors within the bone marrow. A) Flow cytometry gating strategy to identify lineage^-^ (Lin^−^), mature effector cells, hematopoietic stem cells (HSC), and common lymphoid progenitors (CLP) in 21- to 35-day-old mice. B) Proportion of Lin^−^ and mature effector cell populations out of total live cells, identified based on the absence or presence of Gr-1, B220, CD2, CD3, CD5, CD8, and Ter-119 cell surface markers, respectively. C) Proportion of HSC (Lin^−^Sca-1^+^c-Kit^+^CD150^+^CD48^−^) and CLP (Lin^−^Sca-1^+^c-Kit^+^IL7R- α^+^/CD127^+^) populations out of total live cells. Each point in B and C represents an individual mouse. D) Total number of CFUs present per 2x10^4^ bone marrow cells after 10 days of growth on complete methylcellulose media. Each point represents an independent culture, and duplicate cultures were plated for each mouse. For all panels, *n* ≥ 5 mice per genotype, mean ± SEM are plotted, and significance was determined via unpaired *t*-test (\**P*≤0.05, \*\**P*≤0.01, \*\*\**P*≤0.001, and \*\*\*\**P*≤0.0001).

Based on our observations of marked anemia and overall reduction in total bone marrow cells, thymocytes, mature effector cells, and circulating WBCs, but not HSCs, we hypothesized that reduced MRN may impact survival of HSPCs as they undergo differentiation and proliferation. To address this question, we examined the survival of bone marrow progenitors via the colony formation assay and found that *TgWT(L)* and *TgATLD1(L)* exhibited clear defects in the ability to form colonies compared to controls (Fig. 3D). Together, these findings demonstrate that reduced expression of wildtype MRE11 or MRE11-ATLD1 significantly impairs HSPC differentiation and proliferation. However, hematopoiesis was not significantly impacted in *TgATLD1(H)*, indicating the MRE11 C-terminus is not essential for the development of blood cell progenitors in the bone marrow.

### Impaired lymphocyte development in mice expressing reduced MRN

The bone marrow and thymus are the primary sites of postnatal B and T lymphocyte development, respectively. Based on our observations of markedly reduced numbers of live cells within these lymphoid organs in *TgWT(L)* and *TgATLD1(L)* mice (Fig. 2), we hypothesized that reduced MRN and/or loss of the MRE11 C-terminus may impair lymphocyte development. To address this question, we assessed the B and T lymphocyte populations in the *Mre11* transgenic mice by flow cytometry (Fig. 4). In the bone marrow, early pro-B cell progenitors express surface B220 and CD43, then progress to more mature stages upon successful assembly of immunoglobulin heavy (IgH) and light (IgL) chain variable region genes via the site-specific DNA rearrangement, V(D)J recombination. We examined pro-B (B220^+^CD43^+^) and pre-B cell (B220^+^CD43^−^) populations, as well as the immature (B220^lo^IgM^+^) and mature (B220^hi^IgM^+^) B cells, which express surface immunoglobulin M (IgM). In *TgWT(L)* and *TgATLD1(L)* bone marrow, no differences in the percentages of pro-B cells were observed (Fig. 4A). However, we observed significantly reduced percentages of pre-B, immature, and mature B cell populations (Fig. 4A, B). In *TgATLD1(H)* bone marrow, the immature and mature B cells populations were also modestly decreased compared to controls (Fig. 4B). These findings indicate that reduced levels of wildtype MRE11 or MRE11-ATLD1 specifically impair early B cell development at the pro-B to pre-B cell transition, which coincides with the initiation of V(D)J rearrangements. The results also demonstrate that loss of the MRE11 C-terminus alone impedes the differentiation of pre-B cells, which clonally proliferate and undergo IgL rearrangements (30), thereby leading to a reduction of immature and mature B cell populations.

**Figure 4.**
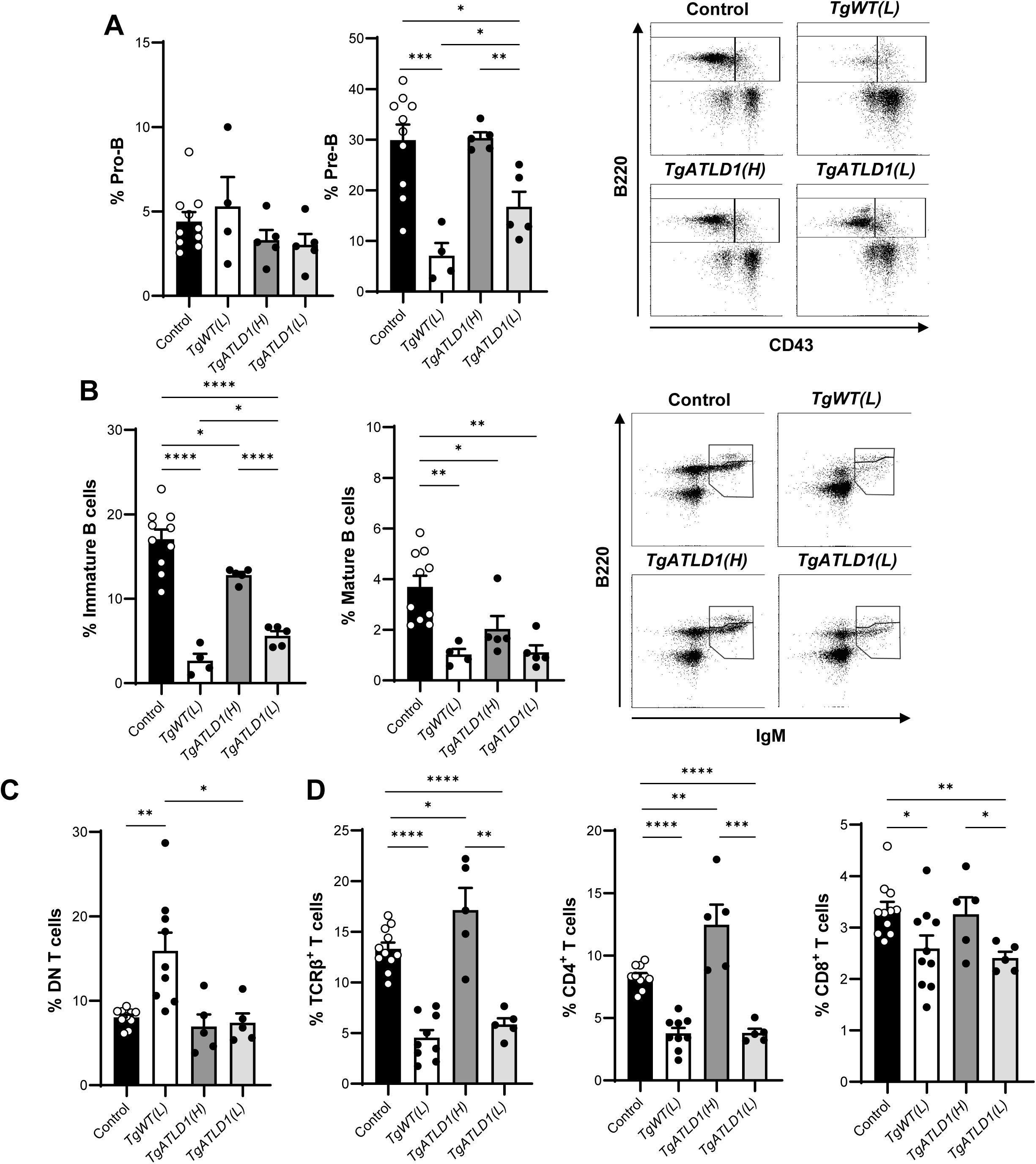
Distinct lymphocyte development defects in *TgATLD1(H)*, *TgATLD1(L)* and *TgWT(L)* mice. A) (Left) Proportion of pro-B (B220^+^CD43^+^) and pre-B (B220^+^CD43^−^) cell populations out of total lymphocytes in the bone marrow. (Right) Representative B220 versus CD43 flow cytometry plots for each genotype are shown with the pre-B (left) and pro-B (right) square gates drawn. B) (Left) Proportion of immature (B220^+^IgM^low^) and mature (B220^+^ IgM^high^) B cells out of total lymphocytes in the bone marrow. (Right) Representative B220 versus IgM flow cytometry plots for each genotype are shown with the immature (bottom) and mature (top) polygon gates drawn. C) Proportion of DN (CD4^−^CD8^−^), TCRβ^+^, CD8^+^, and CD4^+^ T cells out of total lymphocytes in the thymus. For all panels, each point represents an individual mouse, *n* ≥ 5 mice per genotype, mean ± SEM are plotted, and significance was determined via unpaired *t*-test (\**P*≤0.05, \*\**P*≤0.01, \*\*\**P*≤0.001, and \*\*\*\**P*≤0.0001).

During T cell development in the thymus, CD4^−^CD8^−^ double negative (DN) progenitors transition to CD4^+^CD8^+^ double positive (DP) then CD4^+^ or CD8^+^ single positive (SP) T cells that express surface TCRβ. We observed a significant accumulation of DN progenitors in *TgWT(L)* mice and a concomitant decrease in TCRβ^+^ T cells in *TgWT(L)* mice (Fig. 4C, D). In *TgATLD1(L)* mice, the percentage of DN progenitors was indistinguishable from wildtype controls, but similar to *TgWT(L)* mice, the percentage of TCRβ^+^ cells was significantly reduced (Fig. 4C, D). These findings indicate that T cell development is impaired at the stage when cells undergo V(D)J recombination within the TCRβ locus in *TgWT(L)* and *TgATLD1(L)* mice. We observed that CD4^+^ and CD8^+^ single positive (SP) thymocyte populations were also decreased in *TgWT(L)* and *TgATLD1(L)* thymuses, indicating the DP to SP transition, which is dependent on a diverse TCRβ repertoire for positive selection, is also impaired (Fig. 4D). In comparison, T cell development at the stages examined was largely unaffected in *TgATLD1(H)* thymuses. Together, these findings demonstrate that reduced expression of wildtype MRE11 or MRE11-ATLD1 results in defective early B and T cell development at stages when V(D)J rearrangements are initiated. The results also demonstrate that the MRE11 C-terminus is largely dispensable during early stages of lymphocyte development. However, the reduced immature and mature B cell populations observed in *TgATLD1(H)* mice suggest that loss of the MRE11 C-terminus impairs later stages of B cell development which require proliferation and successful rearrangement of IgL loci.

### Splenomegaly associated with extramedullary hematopoiesis, myeloproliferation, and genome instability in mice expressing reduced levels of MRE11 or MRE11-ATLD1

One striking, fully penetrant phenotype observed in *TgWT(L)* and *TgATLD1(L)* mice is significantly enlarged spleens (Fig. 5A). Histologic analyses of hematoxylin and eosin-stained sections reveal a paucity of lymphoid follicles, and red pulp was diffusely expanded with myeloid progenitor cells and scattered plasma blood cells (Fig. 5B). Under conditions of bone marrow insufficiency, HSPCs are mobilized from the bone marrow to extramedullary sites where they can differentiate into effector cells through a process known as extramedullary hematopoiesis (EMH) (31, 32). As the spleen is a primary site of EMH, we examined *TgWT(L)* and *TgATLD1(L)* splenocyte populations by flow cytometry and observed a significant increase in Lin^−^ and LK hematopoietic progenitors with a concomitant decrease in mature effector cells in comparison to controls (Fig. 5C). While *TgWT(L)* spleens also exhibited elevated HSCs and CLPs, consistent with splenic EMH (Fig. 5D), *TgATLD1(L)* spleens did not. This suggests that reduced expression of the MRE11-ATLD1 allele may impact the survival and differentiation of splenic HSPCs at distinct stages compared to *TgWT(L)*. Furthermore, we observed significantly more colonies in *TgWT(L)* and *TgATLD1(L)* spleens than controls via CFU assay (Fig. 5E), confirming elevated hematopoietic progenitor populations. Notably, *TgATLD1(H)* mice did not exhibit EMH phenotypes.

**Figure 5.**
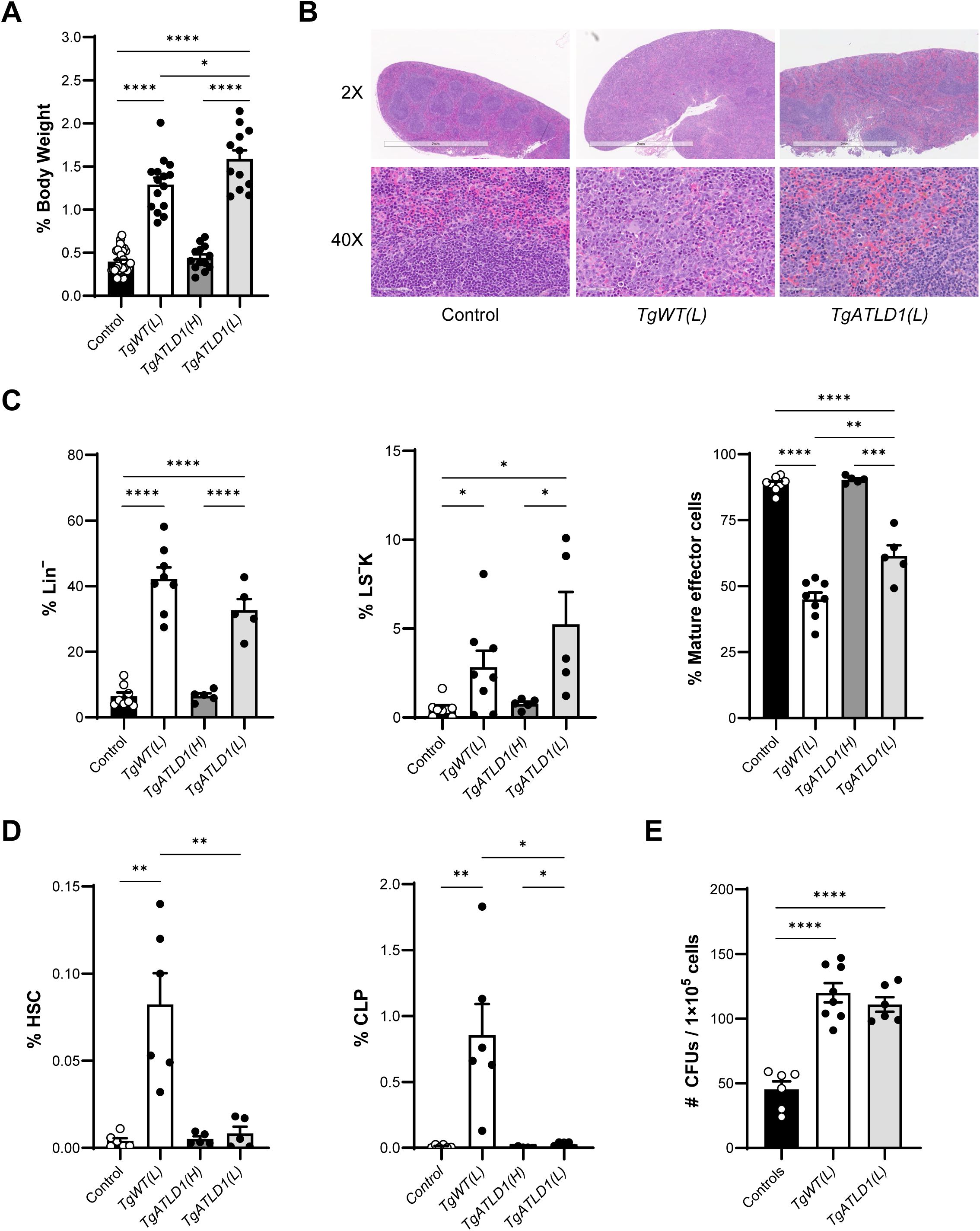
*TgWT(L)* and *TgATLD1(L)* mice exhibit splenomegaly and extramedullary hematopoiesis. A) Spleen weights expressed as a percentage of the total body weight of the mouse. Mice aged 21-100 days were sacrificed and total body weight and splenic weight was determined at time of death. B) Hematoxylin and eosin-stained spleen sections from age-matched 4 week old *TgWT(L)* and *TgATLD1(L)* mice compared to controls. Representative images at two magnifications are shown. C) Percentages of Lin^−^ (Lin^−^Sca-1^−^), LS^−^K ((Lin^−^Sca-1^−^c-kit^+^), and mature effector cell populations out of total live cells. Lin^−^ and mature effector cells were identified based on the absence or presence of Gr-1, B220, CD2, CD3, CD5, CD8, and Ter-119 cell surface markers, respectively. D) Percentages of HSC and CLP (Lin^−^Sca-1^+^c-Kit^+^IL7R-α^+^/CD127^+^) populations out of total live cells. E) CFU assay of splenocytes isolated from mice of the indicated genotypes. Each point represents an independent culture, and duplicate cultures were plated for each mouse. For all panels, *n* ≥ 3 mice per genotype and in all bar plots, each point represents an individual mouse, mean ± SEM are plotted, and significance was determined via unpaired *t*-test (\**P*≤0.05, \*\**P*≤0.01, \*\*\**P*≤0.001, and \*\*\*\**P*≤0.0001).

Based on the spleen histology, we further examined myeloid populations by flow cytometry and found increased percentages of Gr-1^+^Mac-1^+^/CD11b^+^ immature myeloid cells in *TgWT(L)* and *TgATLD1(L)* spleens, consistent with the H&E-stained sections (Fig. 5B) and indicative of a myeloproliferative phenotype (Fig. 6A). Immunohistochemical analyses with the proliferation marker, Ki-67, revealed a significantly higher proportion of Ki-67^+^ nuclei in *TgWT(L)* spleens compared to controls (Suppl. Fig. S2). *TgATLD1(L)* spleens also exhibited consistently higher proportions of Ki-67 positive cells, though the phenotype was less pronounced. Ki-67 was predominantly located within the peripheral red pulp, which is populated by myeloid precursors (Suppl. Fig. S2). Another proliferation marker, the DNA replication protein PCNA, was detected at significantly elevated levels in *TgWT(L)* and *TgATLD1(L)* spleens by western blot (Fig. 6B). Thus, reduced expression of wildtype MRE11 and MRE11-ATLD1, but not loss of the MRE11 C-terminus alone, results in enlarged spleens populated by highly proliferative myeloid lineage progenitor cells.

**Figure 6.**
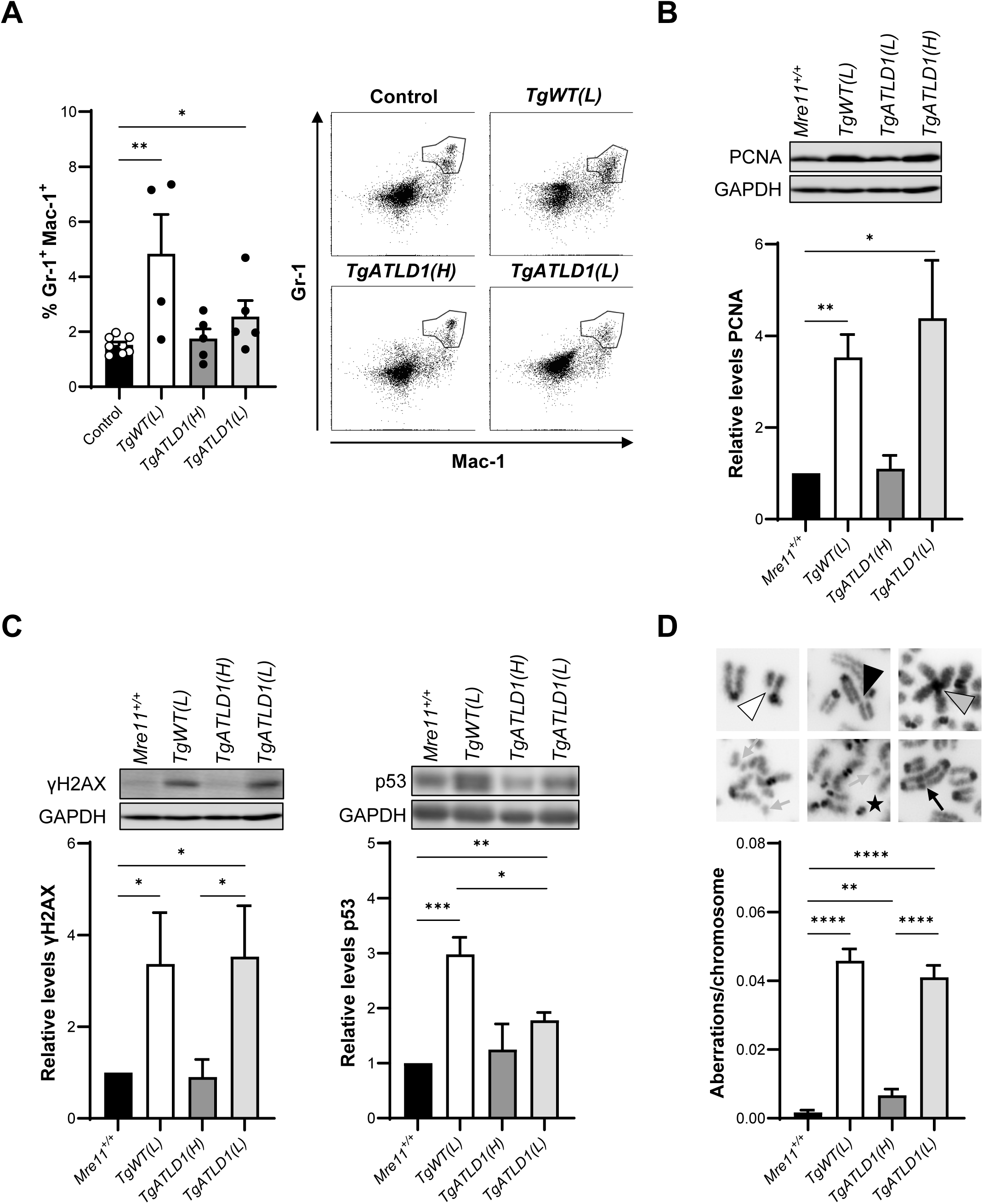
Splenomegaly in *TgWT(L)* and *TgATLD1(L)* mice associated with myeloproliferation and genome instability. A) (Left) Percentages of Gr-1^+^Mac-1^+^/CD11b^+^ immature myeloid cell populations out of total live cells. Cells gated on B220^−^ and CD3^−^ to remove lymphocyte populations. Each point represents an individual mouse for *n* ≥ 4 mice per genotype. (Right) Representative Gr-1 versus Mac-1/CD11b flow cytometry plots for each genotype are shown with the Gr-1^+^Mac-1^+^/CD11b^+^ polygon gate drawn. B and C) Tissue lysates from *TgWT(L)* and *TgATLD1(L)* splenocytes were analyzed by immunoblotting for PCNA (B), γH2AX and p53 (C), GAPDH is the loading control (*n* ≥ 4 animal replicates, aged 5-21 weeks). D) Chromosomal aberrations in metaphase spreads from primary splenocytes isolated from *TgWT(L)*, *TgATDL1(H)* and *TgATLD1(L)* mice were quantified. Chromosome breaks (black arrowhead), chromatid breaks (white arrowhead), Robertsonian translocations (grey arrowhead), chromosome fusions (black arrow), chromosome fragments (grey arrow) and detached centromeres (star). Approximately 40 metaphases from *n* ≥ 3 mice were analyzed. For all bar plots, mean ± SEM is plotted and statistical significance was determined via unpaired *t*-test (\**P*≤0.05, \*\**P*≤0.01, \*\*\**P*≤0.001 and \*\*\*\**P*≤0.0001).

Due to the importance of MRE11 in DSB repair and maintaining genome stability, we examined accumulation of DNA damage in spleens from the *Mre11* transgenic mice. To this end, we performed western blotting for the DNA DSB marker, γH2AX, the phosphorylated form of the histone variant H2AX (phospho-S139). Markedly elevated levels of γH2AX were observed in the enlarged *TgWT(L)* and *TgATLD1(L)* spleens, thereby indicating high levels of unrepaired DSBs (Fig. 6C). We also observed significantly increased p53 levels, an indicator of the DNA damage response. However, p53 levels are lower in *TgATLD1(L)* spleens than *TgWT(L)*, consistent with the reported defect in ATM-dependent DNA damage signaling in ATLD1 cells (1, 19, 26). Increased levels of unrepaired DSBs in hyperproliferating cells can lead to chromosomal instability through failed or aberrant repair of damage. To determine whether *Mre11* transgenic mice exhibit chromosomal instability, we examined the accumulation of chromosomal anomalies in metaphase spreads from primary splenocytes. A striking increase in the number of aberrations in *TgWT(L)* and *TgATLD1(L)* splenocytes was observed (Fig. 6D) and included elevated numbers of chromosome fragments, chromosome and chromatid breaks, fusions, and other complex anomalies (Suppl. Table S1). We found that *TgATLD1(H)* splenocytes also exhibited a significant increase in chromosomal aberrations relative to controls (4-fold), suggesting that the MRE11 C-terminus supports the efficient repair of spontaneous chromosomal DSBs in primary cells. These results are consistent with the previously reported roles of the MRE11 C-terminus in mediating CtIP-dependent repair of DSBs in unperturbed cells and regulating MRE11 DNA binding and nucleolytic activities through post-translational modifications (5, 23–25, 33). Together, these observations suggest that both reduced MRN levels and loss of the MRE11 C-terminus contribute to defects in DSB repair in primary cells and tissues.

### Distinct cellular responses to DNA damage in *Mre11* mutant transgenic cells

The MRN complex is a sensor of DNA ends and upon binding DSBs, recruits the ATM serine/threonine protein kinase. ATM then undergoes autophosphorylation, which activates its intrinsic kinase activity that phosphorylates many downstream protein substrates to elicit cellular responses, including cell cycle arrest, DNA repair, and/or apoptosis. Thus, ATM is considered the master controller of cellular responses to DNA DSBs (34). We previously demonstrated that *TgATLD1(L)* cells exhibit impaired ATM-dependent cellular responses to ionizing radiation (IR)-induced DNA damage. In contrast, *TgWT(L)* cells retained normal ATM-dependent phosphorylation and cell cycle checkpoint responses (26). To assess the importance of the MRE11 C-terminus in activating ATM, IR-induced phosphorylation of the well-characterized ATM substrate KAP1 was assessed in *TgATLD1(H)* MEFs via western blotting (Fig. 7A). As anticipated, pKAP1 (phospho-S824) levels were significantly lower at both 2.5 and 10Gy in *TgATLD1(L)* MEFs compared to wildtype cells, whereas *TgWT(L)* MEFs exhibited similar or higher KAP1 phosphorylation at 2.5 and 10Gy, respectively (Fig. 7A; (26)). We found that *TgATLD1(H)* MEFs exhibited a robust DNA damage response as evidenced by the significant increase in IR-induced pKAP1 that was indistinguishable from wildtype cells (Fig. 7A). Pre-treatment with the ATM kinase inhibitor, KU55933 (ATMi), abolished pKAP1 in all cell lines, thereby demonstrating that phosphorylation is ATM-dependent (Fig. 7A). These findings suggest that the MRE11 C-terminus does participate in efficiently activating ATM; however, near wildtype levels of MRE11-ATLD1 can compensate for the observed defects in ATM activation.

**Figure 7.**
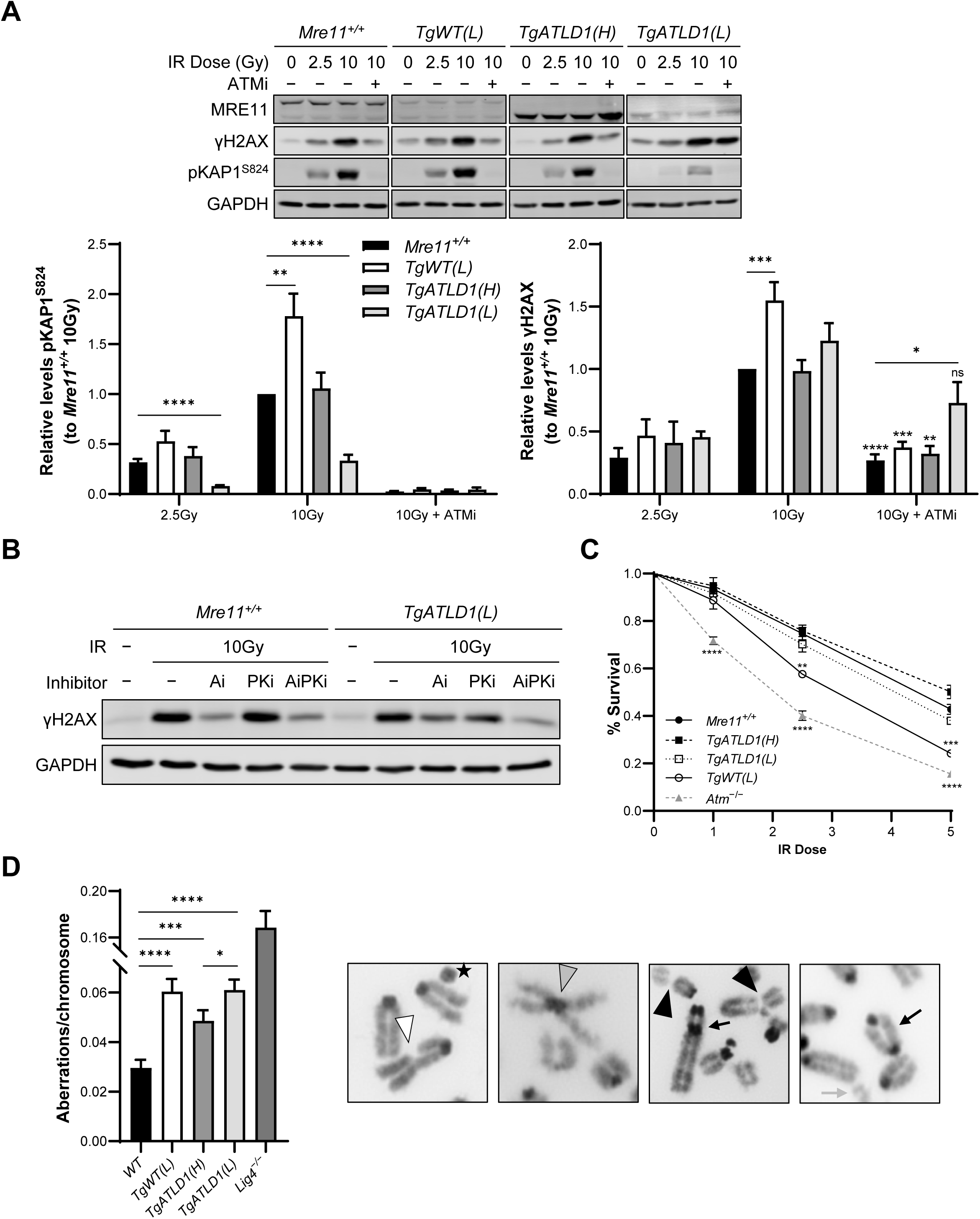
Cells expressing different levels of MRE11 and MRE11-ATLD1 have distinct cellular responses to DNA damage. A) Cell lysates from immortalized MEFs were prepared 30 min after the indicated dose of ionizing radiation and then analyzed by immunoblotting for γH2AX and pKAP1^S824^ levels. Prior to irradiation cells were pretreated with or without the ATM kinase inhibitor KU55933 (ATMi) for 1hr. GAPDH is the loading control. Mean ± SEM is shown (n ≥ 3 cell culture replicates). Right bar graph; asterisks above individual bars in 10Gy + ATMi γH2AX samples indicate significant differences between 10Gy vs. 10Gy + ATMi for each genotype. (ns, not significant, \*\**P*≤0.01, \*\*\**P*≤0.001, and \*\*\*\**P*≤0.0001) B) Cell lysates from immortalized wildtype and *TgATLD1(L)* MEFs were pretreated with the ATM inhibitor (Ai) or a targeted inhibitor of DNA-PKcs (PKi), or both (AiPKi), for 1hr prior to receiving 10Gy of IR. γH2AX levels were assessed by western blotting at 0.5hr post-IR. GAPDH is the loading control. C) MEFs of the indicated genotypes were plated and allowed to grow for 1d prior to exposure to various doses of IR. Cultures were then allowed to grow for 3d post-IR before surviving cell numbers were quantitated. Data are expressed as the percent survival relative to the unirradiated control of the same genotype. *n* ≥ 4 cell culture replicates. D) The number of chromosomal aberrations in metaphase spreads of irradiated immortalized *TgWT(L)*, *TgATLD1(H)* and *TgATLD1(L)* MEFs were quantified. Cells were plated and allowed to grow for 48hr prior to treatment with 2.5Gy IR. Following IR treatment cells were allowed to recover for 24hr post-IR before addition of colcemid and preparation of metaphase spreads. Approximately 40 metaphases from *n* ≥ 3 cell culture replicates were analyzed. Representative images of different chromosomal anomalies. Chromosome breaks (black arrowhead), chromatid breaks (white arrowhead), Robertsonian translocations (grey arrowhead), chromosome fusions (black arrow), chromosome fragments (grey arrow) and detached centromeres (star). In all graphs mean ± SEM are plotted, and significance was determined via unpaired *t*-test (\**P*≤0.05, \*\**P*≤0.01, \*\*\**P*≤0.001, and \*\*\*\**P*≤0.0001).

In addition to KAP1, activated ATM phosphorylates H2AX in response to DSBs. A related protein kinase, DNA-PKcs, is also activated by DSBs upon interaction with the KU70/KU80 DNA end binding complex (known as the DNA-PK holoenzyme), which primarily functions within the NHEJ pathway. In wildtype cells, ATM is the primary kinase responsible for H2AX phosphorylation; however, in cells deficient for MRE11 (*Mre11^−/−^*), DNA-PK can efficiently compensate for ATM to generate γH2AX in response to DSBs (35). Therefore, we examined the relative contributions of ATM and DNA-PKcs to H2AX phosphorylation in *TgATLD1(L)* cells. To this end, cells were pre-treated with ATMi, a targeted inhibitor of DNA-PKcs (NU7026, PKi), or both, and γH2AX levels were assessed by western blotting at 0.5hr post 10Gy IR. ATM inhibition markedly reduced γH2AX in wildtype, *TgWT(L),* and *TgATLD1(H)* cells. However, γH2AX levels were not significantly affected in *TgATLD1(L)* cells, indicating robust phosphorylation occurred independently of ATM (Fig. 7A, B). Exposure of wildtype cells to PKi did not have a measurable impact on H2AX phosphorylation post-IR, and treatment with both inhibitors did not decrease γH2AX beyond ATM inhibition alone (Fig. 7B). Similar results were observed in *TgWT(L)* and *TgATLD1(H)* cells with intact ATM signaling (Suppl. Fig. S3A). In contrast, inhibition of DNA-PKcs significantly reduced H2AX phosphorylation in *TgATLD1(L)* cells, and addition of both ATMi and PKi reduced γH2AX to levels below either alone (Fig. 7B). These findings suggest that, similar to *Mre11^−/−^* cells which completely lack MRN, reduced expression of MRE11-ATLD1 permits KU-dependent activation of DNA-PKcs, thereby compensating for impaired ATM activity for IR-induced H2AX phosphorylation. This mechanism is specific to loss of the C-terminus, as low expression of wildtype MRE11 supports efficient ATM-dependent DNA damage responses (Fig. 7A; (26)).

We next assessed the DNA DSB repair capacity of *TgATLD1(H)* MEFs compared to *TgWT(L)* and *TgATLD1(L)* cells by quantitating relative cellular survival upon exposure to increasing doses of IR (Fig. 7C). As previously reported, *TgWT(L)* MEFs exhibited reduced survival upon exposure to IR but were less sensitive in comparison to *Atm^−/−^* MEFs (26). In contrast, *TgATLD1(H)* MEFs exhibited similar survival at all doses of IR compared to wildtype controls (Fig. 7C), similar to *TgATLD1(L)* MEFs. We also examined the accumulation of chromosomal anomalies in irradiated cells to further assess the repair efficiency of induced DSBs in the *Mre11* transgenic MEFs. To this end, *TgATLD1(H)*, *TgATLD1(L)*, *TgWT(L)*, and controls were exposed to 2.5Gy of IR, and the number and types of chromosomal anomalies were quantitated at 24hr post-IR in a blinded manner. *TgWT(L)*, *TgATLD1(L)*, and *TgATLD1(H)* MEFs all exhibited significantly higher levels of chromosomal anomalies compared to controls (Fig. 7D). Together, these findings indicate that both reduced MRE11 expression levels and loss of the MRE11 C-terminus impair the efficient repair of IR-induced DSBs. However, cells are permitted to survive with elevated chromosomal damage when MRE11-ATLD1 expression is reduced below a threshold level.

We previously observed that *TgATLD1(L)* MEFs exhibit a defective IR-induced G2/M cell cycle checkpoint, which could allow cells harboring chromosomal breaks to progress through the cell cycle and permit DSB repair by alternative pathways (26). Thus, we examined G2/M checkpoint activation in *TgATLD1(H)* MEFs by measuring the mitotic index as determined by percentage of cells positive for the mitosis-entry marker phospho-histone H3^S10^ after exposure to 10Gy IR (36). We observed that the mitotic index in *TgATLD1(H)* MEFs was indistinguishable from wildtype controls (Suppl. Fig. S3B). In combination with the moderate elevation of chromosomal anomalies observed in *TgATLD1(H)* splenocytes, these findings suggest that while loss of the MRE11 C-terminus impairs DNA DSB repair, expression of MRE11-ATLD1 at near normal levels retains sufficient activity to activate the G2/M cell cycle checkpoint.

## Discussion

In this study, we examined the *in vivo* importance of the MRE11 C-terminus in a novel transgenic mouse model expressing the *Mre11-ATLD1* disease allele at near endogenous levels in an *Mre11^−/−^* background. We provide evidence that loss of the MRE11 C-terminus impairs B cell development in the bone marrow at the pre-B to immature B cell transition and results in accumulation of chromosomal damage in primary cells and irradiated transformed cell lines. These findings demonstrate that the MRE11 C-terminus has *in vivo* functions and facilitates efficient repair of chromosomal DSBs. Surprisingly, *TgATLD1(H)* mice did not exhibit severe phenotypes affecting other organ systems and DNA damage responses. In sharp contrast, we observed that significantly reduced expression of either MRE11-ATLD1 or wildtype MRE11 causes marked phenotypes, including bone marrow failure, impaired lymphocyte development, extramedullary hematopoiesis, and splenomegaly. We also found that reduced MRE11-ATLD1 levels result in cellular and organismal phenotypes distinct from low expression of wildtype MRE11, thereby indicating that loss of the MRE11 C-terminus causes a broader range of phenotypes when expression is reduced below a threshold level.

Previous studies of an *Mre11* conditional knock-out mouse model from the Ferguson lab demonstrated that MRE11 is required for early embryonic development, whereas *Mre11* heterozygous mice and cells do not exhibit observable phenotypes (37). In the current study, we demonstrate that reduction of wildtype MRE11 and MRE11-ATLD1 to approximately 10-40% of endogenous MRE11 expression uncovered a critical role for MRN in hematopoiesis, as evidenced by an accumulation of lineage negative progenitors and decrease in mature effector cells within the bone marrow and marked impairment of HSPCs to form colonies in the CFU assay (Figs. 1,2). It is notable that rare *NBS1* and *RAD50* variants cause genome instability syndromes that can also feature severe anemia (17, 28, 38, 39). Recent work has shown that a rapid surge of transcriptional reprogramming during hematopoiesis generates high levels of DNA damage associated with accumulation of nuclear aldehydes and DNA interstrand crosslinks (ICLs) (40). We previously reported that *TgWT(L)* cells are hypersensitive to ICLs, raising the possibility that impaired MRN function and/or low MRN levels may decrease ICL repair capacity and impact the survival of developing blood cells (26). Although hematopoietic defects have not yet been identified in ATLD1 patients, our findings suggest that variants that reduce MRE11 expression, including those that do not significantly affect MRE11 activities, may cause bone marrow failure in patients and have clinical relevance for diagnosis and treatment of these phenotypes.

Bone marrow failure and severe anemia can promote extramedullary hematopoiesis (EMH) that manifests as splenomegaly (31). Indeed, one striking phenotype observed in the *TgWT(L)* and *TgATLD1(L)* mice was significantly enlarged spleens (Fig. 5A) harboring lineage negative hematopoietic progenitor cells, which were largely absent from control spleens (Fig. 5C). The spleens were also dysmorphic with expanded red pulp that contained myeloid progenitor cells and stained positively for the proliferation marker, Ki-67. Splenocytes isolated from the *TgWT(L)* and *TgATLD1(L)* spleens were also characterized by elevated PCNA levels and significant genomic damage, including accumulation of chromosome and chromatid breaks and fusions.

EMH can lead to tumorigenesis due to production of myeloid lineage cells that promote tumor growth (31, 41). Furthermore, genome instability syndromes characterized by bone marrow failure, such as Fanconi anemia, strongly predispose to malignancies (29, 42). Our observations that *TgWT(L)* and *TgATLD1(L)* spleens harbor highly proliferative myeloid progenitors and accumulate chromosome anomalies suggest reduced MRN levels may predispose to myeloproliferative neoplasms. However, we have not yet observed increased incidence of spontaneous cancers or solid tumors arising in the *TgWT(L)* or *TgATLD1(L)* transgenic mice. It is possible that these transgenic mice may exhibit cancer predisposition if they are bred onto a background that promotes tumorigenesis, such as p53-deficiency, to accelerate cancer initiation and progression.

Our analyses revealed that *TgWT(L)* and *TgATLD1(L)* mice exhibit defective B and T lymphocyte development at the specific stages when V(D)J rearrangements are initiated. We also observed defects in the pre-B to immature B cell transition in *TgATLD1(H)* mice, the stage when Ig light chain rearrangements occur to produce mature B cell receptors. V(D)J recombination is initiated by the RAG1/2 endonuclease which cleaves DNA at recombination signal sequences adjacent to V, D, and J gene segments. This cleavage generates DSBs with two distinct end structures: covalently closed, hairpin coding ends, and blunt, 5’-phosphorylated signal ends. RAG1/2 sequesters the ends in a post-cleavage complex to ensure coordinated end processing and joining via the classical NHEJ pathway. MRN localizes to RAG1/2-induced DSBs and is hypothesized to stabilize post-cleavage complexes (43–45). Moreover, defects in MRN function promote aberrant transrearrangements involving V(D)J loci on different chromosomes. These findings have led to the hypothesis that MRN functions during V(D)J recombination to promote normal rearrangements and/or suppress aberrant interchromosomal events (19, 45–47). Our results in the *Mre11* transgenic mice now provide evidence that MRN facilitates normal lymphocyte development. It will be of interest to examine localization of MRE11-ATLD1 to V(D)J loci as well as assess the frequency of normal and aberrant V(D)J recombination and sequences of V(D)J junctions in *TgWT(L)*, *TgATLD1(L)* and *TgATLD1(H)* to define *in vivo* roles for MRN and the MRE11 C-terminus during V(D)J recombination.

Experiments in *TgATLD1(L)* MEFs revealed that ATM-dependent DNA damage responses are impaired and, unlike wildtype MEFs, DNA-PKcs rather than ATM is the primary kinase that phosphorylates H2AX (Fig. 7B). These results parallel those observed in *Mre11^−/−^*MEFs that completely lack MRN, which exhibit defective ATM activation and phosphorylation of H2AX in a KU- and DNA-PKcs-dependent manner, indicating that the DNA-PK holoenzyme occupies DNA ends that would otherwise be bound by MRN to elicit cellular responses to DSBs (35). As ATM activation is not defective in *TgWT(L)* MEFs, these findings suggest that loss of the MRE11 C-terminus in conjunction with reduced expression levels may impair efficient MRN localization to chromosomal DSBs leading to compensatory DNA-PKcs activation by KU. In this regard, the MRE11 C-terminus binds intact double-strand DNA independently of its end-binding activity (5, 48) and is hypothesized to be required for genome scanning to detect and remove DNA-PK bound to DNA ends (49). Further investigation into the efficiency of MRE11-ATLD1 versus KU localization to chromosomal DSBs could reveal important mechanistic insights into the interplay between MRN and KU *in vivo* and provide insights into the functional importance of the MRE11 C-terminus during general DSB repair.

We did not observe DNA damage response defects, IR hypersensitivity, or impaired cell cycle defects in *TgATLD1(H)* transformed MEFs (Fig. 7A, C; Supp. Fig. 3). These results were unexpected given the functions of the MRE11 C-terminus in DSB repair processes and the distinct cellular phenotypes observed in *TgATLD1(L)* versus *TgWT(L)* MEFs. It is possible that the levels of MRE11-ATLD1 in *TgATLD1(H)* cells and tissues, which are 1.5-2-fold higher than endogenous MRE11, may be high enough to compensate for loss of specific MRN functions. It is worth noting that *TgATLD1(H)* primary splenocytes and irradiated MEFs exhibit a similar significant increase in chromosomal aberrations compared to *TgATLD1(L)* and *TgWT(L)* MEFs, suggesting that the levels of MRE11-ATLD1 may have reached a threshold that can complement a subset of MRN functions, but do not completely compensate for the DNA DSB repair activity. These findings underline the importance of defining MRE11 and MRE11-ATLD1 expression levels that are necessary to support distinct MRN functions and prevent disease-associated phenotypes.

Our study has revealed that the constellation and severity of phenotypes caused by disease-associated *MRE11* variants are significantly influenced by expression levels. We provide evidence that the MRE11 C-terminus, deleted in the *Mre11-ATLD1* allele, does have *in vivo* functions during B cell development and in promoting efficient repair of chromosomal DSBs; however, unexpectedly, it is not essential for the critical roles of MRE11 in DNA damage responses and repair when expression of the truncated protein is enforced. We demonstrate that reduced expression of MRE11-ATLD1 or wildtype MRE11 leads to impaired hematopoiesis, defects in lymphocyte development, and splenomegaly associated with highly proliferative myeloid progenitors. These phenotypes have not been previously attributed to impaired MRE11 function and thus, have important implications for clinical predictions in ATLD patients harboring pathogenic variants that may affect not only specific MRE11 activities, but also expression levels of MRE11 and the MRN complex. Additionally, our findings identify new areas of biology for MRE11 and MRN function in differentiation and survival of HSPCs, V(D)J recombination, and lymphocyte development that warrant further investigation.

## Materials and Methods

### Generation of Transgenic *Mre11* mice

The transgene was constructed in plasmid pCAGEN, a gift from Connie Cepko (Addgene plasmid #11160; http://n2t.net/addgene:11160; RRID:Addgene 11160). Mm*Mre11a* was amplified from pEF6-Mre11 or pEF6-Mre11-ATLD1 (Regal et al., 2013), with the addition of a 5’ EcoRI and 3’ EcoRV site for cloning into pCAGEN. The transgenes were isolated from pCAGEN-Mm*Mre11a* with HindIII and SpeI. Purified DNA was microinjected into fertilized eggs obtained by mating (C57BL/6J X SJL/J)F1 (Jackson Laboratory Stock number 100012) female mice with (C57BL/6J X SJL/J)F1 male mice. Pronuclear microinjection and embryo transfer was performed as described (Becker et al., 2011). The integration sites of the transgenes were identified by whole genome sequencing. All integrants used in this study are in intergenic regions and are at least 0.75Mb away from the nearest gene with known functions in DNA repair.

### Cell culture

MEFs were isolated from day e13.5 embryos and grown in standard culture conditions as described (Hogan et al., 1994). Primary MEFs were immortalized by transfection with pBsSVD2005 (SV40 large T antigen expression vector). Where indicated, cells were treated with ionizing radiation (^137^Cs source). When indicated, cells were pre-treated with the indicated kinase inhibitor (ATM inhibitor, KU55933, Cayman Chemical; DNA-PK inhibitor, NU7026, Cayman Chemical) at 10μM (Ai) or 20uM (PKi) for 1hr prior to IR (^137^Cs source).

Unless otherwise noted, cells were cultured in DMEM (Gibco) containing 4.5g/L glucose, 1mM sodium pyruvate, 6mM L-glutamine, 100U/mL penicillin, 100μg/mL streptomycin, 20mM HEPES, 1X non-essential amino acids, 90μM 2-mercaptoethanol and 10% heat-inactivated FBS. All cells were grown in a humidified chamber at 37°C containing 5% CO2.

### Determination of Mendelian Inheritance

Mice are *Mus musculus* of C57B6/129sv mixed background. Chi-squared (Χ^2^) analysis was used to determine the significance of any observed differences between the actual and expected results of the crosses performed.

### PCR-based Genotyping

The PCR conditions used to distinguish endogenous *Mre11* alleles were as previously described (Buis et al., 2008). The PCR primers used to identify *TgWT(L)* transgene positive animals are as follows:

1. 5‘-GTG TGG AGG CTG CTT TGA AA-3’
2. 5’-TCA TGT TCA TGG CCC CAG AT-3’
3. 5’-AAC CAC TTG AAT GGG CTC AA-3’

The unique band is 648bp for the transgene. Thermocycling conditions for the above reaction entails 35 cycles of 94°C for 45 sec, 55°C for 45 sec, 72°C for 75 sec.

The PCR primers used to identify *TgATLD1(H)* or *TgATLD1(L)* transgene positive animals are as follows:

1. 5‘-GCC ACC TCA ATC AAC ATC TAG AAA-3’
2. 5’-TGA TGA CCT CTT TAT AGC CAC CT-3’

The unique band is 404bp for the transgene. Thermocycling conditions for the above reaction entails 30 cycles of 94°C for 45 sec, 55.5°C for 45 sec, 72°C for 30 sec.

### Paraffin Processing and Sectioning of Tissues

Briefly, formalin-fixed tissues were processed through graded alcohols and cleared with xylene followed by infiltration with molten paraffin using an automated VIP5 or VIP6 tissue processor (TissueTek, Sakura-Americas). Following paraffin embedding using a Histostar Embedding Station (ThermoScientific), tissues were then sectioned on a rotary microtome (M355S by ThermoFisher Scientific or RM2255 by Leica Biosystems) at 4μm thickness, mounted on glass slides and heated to 60°C for 1 hour.

### Hematoxylin and Eosin Staining of Histologic Tissue Sections

Following deparaffinization and hydration with xylene and graded alcohols, formalin-fixed, paraffin embedded (FFPE) slides were stained with Harris hematoxylin (ThermoFisher Scientific), differentiated with Clarifier (ThermoScientific), blued with bluing reagent (ThermoFisher Scientific), stained with eosin Y, alcoholic (ThermoFisher Scientific), then dehydrated and cleared through graded alcohols and xylene and coverslipped with Micromount (Leica) using a Leica CV5030 automatic coverslipper.

### Ki-67 Staining of Histologic Tissue Sections

All products used are from Biocare Medical (Pacheco, CA) unless otherwise stated. Slides were deparaffinized and rehydrated through a series of xylene, graded alcohols, and distilled water. Deparaffinized slides then underwent heat-induced epitope retrieval (HIER) in a pressure chamber (Decloaking Chamber) for 40 minutes at a maximum temperature of 127°C in a pH 6.0 citrate-based buffer (Diva Decloaker, DV2004). IHC staining was performed on an automated stainer (IntelliPATH FLX) which included steps to quench endogenous peroxidases and block non-specific sites. Anti-mouse rabbit monoclonal Ki67 antibody (ab16667, Abcam) was applied at a 1:750 dilution for 1 hour at 25°C. A negative control slide using naïve mouse/rabbit serum (NC498) in place of primary antibody was run concurrently. Detection was performed using a commercial polymer-based, biotin-free system (IPK5010G80). A DAB-enhancer was applied for 1 minute (DS830). Finally, samples were counterstained with hematoxylin (CATHE) for 5 min, dehydrated and cleared through graded alcohols and xylene, and coverslipped.

### Peripheral Blood Analysis

Blood from sacrificed mice was collected by cardiac puncture immediately following secondary euthanasia. To prevent coagulation, blood was placed into tubes coated with EDTA (Sarstedt). Complete blood counts (CBC) were analyzed using a Hemavet 950 hematology analyzer (Drew Scientific).

### Immunoblotting

Cell extracts were prepared in SDS lysis buffer (10% glycerol; 2% SDS; 62.5mM Tris-HCl, pH 6.8), boiled for 5min, sonicated and cleared by centrifugation (20000 x *g*). Protein concentration was determined by BCA assay (Pierce), β-mercaptoethanol was added to reduce samples, extracts were resolved by SDS-PAGE and transferred using standard procedures.

Images were captured using an Odyssey CLx Imager (LI-COR) and quantification of all immunoblots was done using Image Studio™ Lite Software (LI-COR). Briefly, the signal intensity for the target protein was corrected for protein loading by dividing it by the signal intensity of the loading control. Next, the relative protein level of the target protein was determined by dividing the corrected signal intensity of the target protein by the corrected signal intensity of the reference protein (resulting in a relative protein level of 1 for the reference protein).

Flash frozen tissues were lysed and homogenized in SDS lysis buffer using a mortar and pestle. Samples were then boiled, sonicated, and cleared by centrifugation (20,000 x *g*). Protein concentration was determined by BCA assay (Pierce), β-mercaptoethanol was added to reduce samples, extracts were resolved by SDS-PAGE and transferred using standard procedures. Antibodies used are listed in Supplemental Methods.

### Colony forming unit assay

Single-cell suspensions of bone marrow cells or splenocytes were prepared and then treated with ACK Lysing buffer (Gibco). Cells were then washed in PBS with 5% FBS and then resuspended in IMDM with 2% FBS. The number of viable cells was determined via trypan blue staining and hemocytometer counting. Cells were then diluted to a concentration of 2x10^5^ cells/mL (BM) or 1x10^6^ cells/mL (splenocytes) and plated in duplicate in complete MethoCult (M3434, StemCell Technologies) media according to manufacturer’s instructions. Visible colonies were counted by microscopy at day 10 post-plating.

### Hematopoietic Cell Analyses and Flow Cytometry

Hematopoietic progenitors, myeloid, and lymphocyte populations were analyzed by flow cytometry using single-cell suspensions of bone marrow, spleen, and thymus from mice aged 27-35 days. Cell suspensions from bone marrow and spleen were treated with ACK Lysing buffer (Gibco) to deplete red blood cells. All suspensions were maintained in PBS with 5% FBS. Data was collected using a Novocyte Quanteon (Agilent) and analyzed using FlowJo version 10 software. Labeled antibodies and gating strategies used are listed in Supplemental Methods.

### IR Sensitivity Assays

For IR hypersensitivity assays, MEFs were plated in a 24-well dish at a low density (2500-4000 cells/well) and allowed to attach overnight. Cells were then treated with varying doses of IR and allowed to grow until the unirradiated control well for each genotype reached confluency (3 to 4 days). Cellular survival was determined using a crystal violet colorimetric assay, as previously described (50). Average percent survival was obtained from at least three independent experiments.

### Metaphase Chromosome Spreads and Analysis

Metaphase spreads were generated from early passage cultured immortalized mouse embryonic fibroblasts (MEFs) or primary murine splenocytes harvested from mice between 26 and 33 days of age. MEFs were irradiated with 2.5Gy IR (^137^Cs) and allowed to recover for 24 hours. Irradiated MEFs and primary unirradiated splenocytes were treated with 200 mg/mL colcemid for 3 hours, washed once in PBS, resuspended in 0.4% KCl, and fixed with a 3:1 methanol:glacial acetic acid solution. Fixed cells were dropped onto slides and stained with DAPI. At least 40 metaphase images per culture (MEFs) or mouse (splenocytes) were analyzed for *n* ≥ 3 independent cell culture replicates or animals. Images were blinded before analysis.

#### G2/M Checkpoint Assay

5x10^5^ MEFs were plated per 10cm dish, grown for 48 hr, treated with or without 10Gy IR (^137^Cs), allowed to recover for 1 hour and fixed. Cells were analyzed for the mitotic marker phospho-histone H3^S10^ (Cell Signaling, #9706) (36) and FITC-conjugated secondary antibody (BD Biosciences). Flow cytometry was carried out on a Novocyte Quanteon (Agilent) and analyzed using FlowJo version 10 software as previously described (19).

### Statistics

Statistical analyses were performed using GraphPad Prism software, version 10, with *P* less than 0.05 considered statistically significant. The data are presented as mean ± SEM. The statistical test applied is noted for each experiment.

### Study Approval

All mice were housed at the Taubman Biomedical Science Research Building, University of Michigan (UM). All vertebrate animal experiments were approved by and performed in accordance with the regulations of the University Committee on Use and Care of Animals.

## Acknowledgements

This work was supported by National Institutes of Health, USA grant R01HL15306801 (J.M.S. and D.O.F.) and The University of Michigan Cancer Center Support Grant P30CA046592 to support use of the University of Michigan Transgenic Animal Model Core. M.B.D. received support from the Program in Biomedical Sciences Ph.D. program, University of Michigan Medical School. We thank the University of Michigan Unit for Laboratory Animal Medicine Pathology Core for their immunohistochemistry expertise and diagnostic support.

