## Supplemental Figures for "Differential expression of a disease-associated *MRE11* variant reveals distinct phenotypic outcomes"

#### Supplemental Figure S1

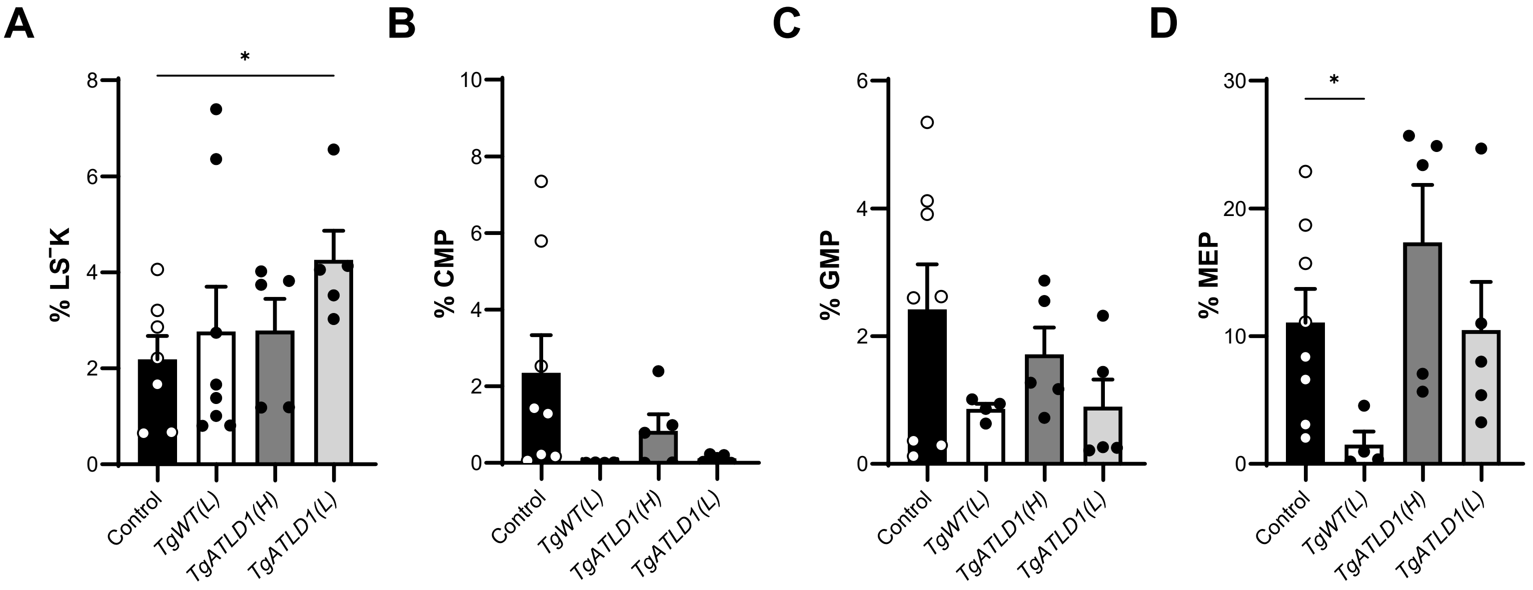


**Supplemental Figure S1. Populations of LS⁻K and myeloid progenitor cell populations were variable in the bone marrow.** Proportion of lineage⁻, Sca-1⁻, and c-Kit⁺ (LS⁻K), common myeloid progenitor (CMP; lineage⁻, Sca-1⁻, c-Kit⁺, CD16/32⁻, CD34⁺), granulocyte-monocyte progenitor (GMP; lineage⁻, Sca-1⁻, c-Kit⁺, CD16/32⁺, CD34⁺), and megakaryocyte-erythroid progenitors (MEP; lineage⁻, Sca-1⁻, c-Kit⁺, CD16/32⁻, CD34⁻). LS⁻K population out of total live cells, CMP, GMP, and MEP populations out of total lineage⁻ cells. Each point represents an individual mouse. $n \geq 4$ mice per genotype. Mean ± SEM plotted. Significance determined via unpaired *t*-test (*$P \leq 0.05$, **$P \leq 0.01$, ***$P \leq 0.001$, ****$P \leq 0.0001$).

### Supplemental Fig. S2

A

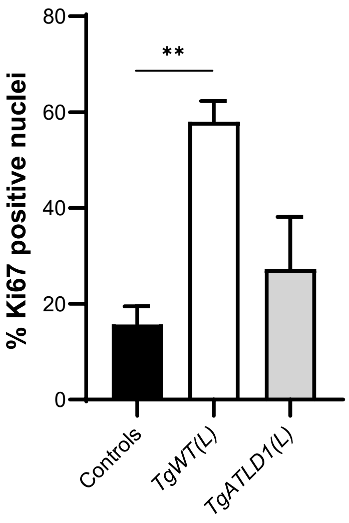


B

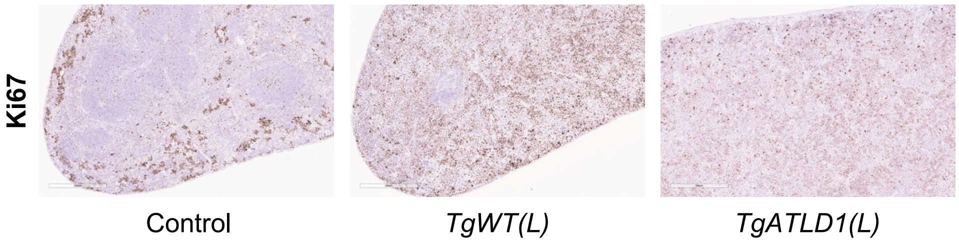


C **Histopathology scores of Ki67-stained spleen sections**

| Genotype | *Mre11*$^{+/+}$ | *TgWT(L)* | *TgATLD1(L)* |
|---|---|---|---|
| Mouse #1 | 1 | 4 | 4 |
| Mouse #2 | 2 | 4 | 4 |
| Mouse #3 | 3 | 4 | − |
| Mean labeling score | 2 | 4 | 4 |

**Supplemental Figure S2. Ki67 staining of spleen sections shows elevated levels of cellular proliferation in mice expression low MRE11.** A) The percentages of Ki67-positive nuclei of the entire stained tissue sections was determined by QuPath analysis (Bankhead, et al. (2017) Sci Rep, 7:16878). B) Representative images of Ki67 stained spleen sections of the indicated genotypes. C) Table of individual Ki67 labeling scores as determined by a University of Michigan veterinarian pathologist using pathology scoring scale of 0 (no labeling), 1 (minimal, <10% of tissue), 2 (mild, 10-25%), 3 (moderate, 25-50%), or 4 (marked nuclear labeling >50%). $n \geq 2$ mice per genotype. Mean ± SEM plotted. Significance determined via unpaired *t*-test (*$P \leq 0.05$, **$P \leq 0.01$).

#### Supplemental Fig. S3

A

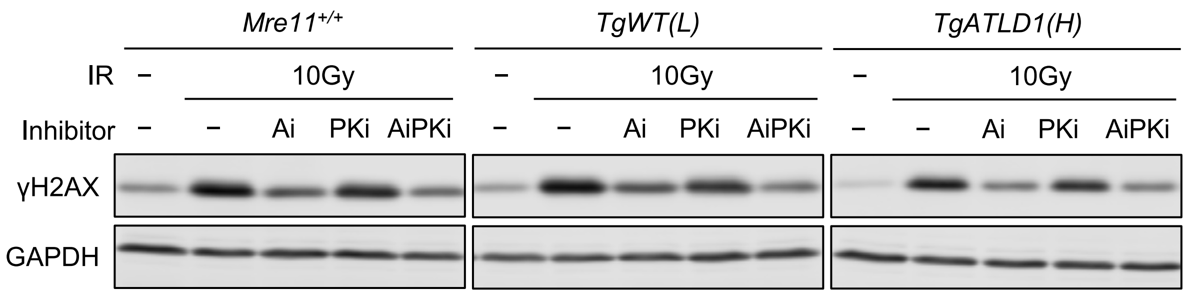


B

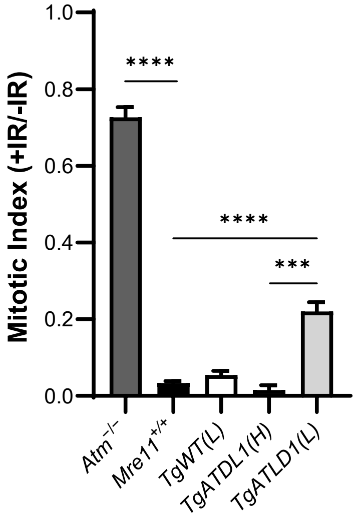


**Supplemental Figure S3. Activation of ATM-dependent DNA damage responses in *Mre11* transgenic MEFs.** A) Cell lysates from immortalized wildtype, *TgWT(L),* and *TgATLD1(H)* MEFs were pretreated with the ATM inhibitor (Ai) or a targeted inhibitor of DNA-PKcs (PKi), or both (AiPKi), for 1hr prior to receiving 10Gy of IR. γH2AX levels were assessed by western blotting at 0.5hr post-IR. GAPDH is the loading control. Representative images are shown. B) Assessment of the G2/M checkpoint by comparison of mitotic indices of MEFs of the indicated genotypes (percentage phospho-$H3^{S10}$-positive cells by flow cytometry) before and 1 hour after 10 Gy IR. Mean ± SEM shown *(n* ≥ 3 cell-culture replicates). Significance determined via unpaired *t*-test (****P*≤0.001, *****P*≤0.0001). The elevated mitotic index in $Atm^{-/-}$ and *TgATLD1(L)* reflects reduced function of the G2/M checkpoint.

### Supplemental Table S1

##### Primary splenocytes

| | *Mre11^+/+^* | *TgWT(L)* | *TgATLD1(H)* | *TgATLD1(L)* |
|---|---|---|---|---|
| Fragments | 0 | 42 | 3 | 66 |
| Detached Centromeres | 0 | 36 | 5 | 31 |
| Chr./Chromatid Breaks | 1 | 100 | 3 | 85 |
| Fusions | 8 | 28 | 4 | 9 |
| Other | 2 | 24 | 13 | 43 |
| Total Chromosomes | 5099 | 4878 | 4232 | 5684 |

##### Irradiated MEFs

| | *Mre11^+/+^* | *TgWT(L)* | *TgATLD1(H)* | *TgATLD1(L)* | *Lig4^–/–^* |
|---|---|---|---|---|---|
| Fragments | 101 | 88 | 158 | 199 | 354 |
| Detached Centromeres | 46 | 45 | 52 | 95 | 166 |
| Chr./Chromatid Breaks | 53 | 68 | 58 | 125 | 260 |
| Fusions | 47 | 84 | 74 | 143 | 250 |
| Other | 57 | 61 | 78 | 149 | 153 |
| Total Aberrations | 304 | 346 | 420 | 711 | 1183 |
| Total Chromosomes | 6467 | 5806 | 10476 | 13202 | 6620 |

**Supplemental Table S1. Breakdown of metaphase aberration types and total chromosomes examined in primary splenocytes and irradiated MEFs.**
