## Supplemental Materials and Methods for "Differential expression of a disease-associated *MRE11* variant reveals distinct phenotypic outcomes"

Primary antibodies used for immunoblotting:

| Antibody | Company | Catalog # |
|---|---|---|
| anti-MRE11 | Cell Signaling Technology | 4895 |
| anti-β-Actin | Proteintech | 66009-1-Ig |
| anti-RAD50 | Bethyl Laboratories | A300-184A |
| anti-NBS1 (Y112) | Novus Biologicals | NB100-57272 |
| anti-p95/NBS1 (Y112) | Abcam | Ab32074 |
| anti-KAP1 (phosS824) | Bethyl Laboratories | A300-767A |
| anti-H2AX (phosS139) | Cell Signaling Technology | 2577 |
| anti-GAPDH (14C10) | Cell Signaling Technology | 2118 |

Secondary antibodies were IRDye conjugated goat anti-rabbit or anti-mouse (LI-COR Biosciences).

Antibodies used for flow cytometry:

| Antibody | Fluorophore | Company | Catalog # |
|---|---|---|---|
| anti-B220 | PE | BioLegend | 103208 |
| anti-CD117 (c-Kit) | APC | BioLegend | 105812 |
| anti-CD11b | PE-Cy7 | eBioscience | 25-0112-82 |
| anti-CD11b | PE | eBioscience | 12-0112-82 |
| anti-CD127 (IL7R-a) | PE-Cy7 | BioLegend | 135013 |
| anti-CD16/CD32 | PE-Cy7 | BioLegend | 101318 |
| anti-CD2 | PE | BioLegend | 100108 |
| anti-CD25 | PE | eBioscience | 12-0251-81 |
| anti-CD3 | PE | BioLegend | 100206 |
| anti-CD34 | FITC | BD Biosciences | 560238 |
| anti-CD3e | FITC | eBioscience | 11-0031-82 |
| anti-CD3e | APC | BioLegend | 100312 |
| anti-CD4 | FITC | eBioscience | 11-0041-82 |
| anti-CD4 | APC | BioLegend | 100412 |
| anti-CD43 | FITC | BD Biosciences | 553270 |
| anti-CD44 | APC | BioLegend | 103011 |
| anti-CD45 | FITC | BioLegend | 103108 |
| anti-CD45R (B220) | FITC | eBioscience | 11-0452-82 |
| anti-CD45R (B220) | APC | eBioscience | 17-0452-82 |
| anti-CD45R (B220) | PerCP-Cy5.5 | eBioscience | 45-0452-82 |
| anti-CD5 | PE | BioLegend | 100608 |
| anti-CD8a | FITC | eBioscience | 11-0081-82 |
| anti-CD8a | PE | BioLegend | 100708 |
| anti-Gr-1 | FITC | BioLegend | 108406 |
| anti-Gr-1 | PE | BioLegend | 108408 |
| anti-IgM | PE | SouthernBiotech | 1021-09 |
| anti-Ly-6A/E (Sca-1) | PerCP-Cy5.5 | BioLegend | 122524 |
| anti-Mac-1 (F4/80) | FITC | BioLegend | 123108 |
| anti-Mac-1 (F4/80) | PE | BioLegend | 123110 |
| anti-pH3 | FITC | BD Biosciences | 553443 |
| anti-TCRβ | FITC | BioLegend | 109206 |
| anti-TCRγδ | FITC | BioLegend | 107504 |
| anti-TER-119 | PE | BioLegend | 116208 |
